# Type I Interferon Signaling Defines a Novel Disease Signature in Xeroderma Pigmentosum C Human Keratinocytes

**DOI:** 10.1101/2025.07.17.665356

**Authors:** Ali Nasrallah, Hamid-Reza Rezvani, Lucid Belmudes, Sandrine Bourgoin-Voillard, Farah Kobaisi, Mohammad Fayyad-Kazan, Michel Seve, Yohann Couté, Eric Sulpice, Walid Rachidi

**Author notes:** Corresponding Authors: Walid Rachidi.

## Abstract

Xeroderma Pigmentosum C (XPC) is a DNA damage recognition protein central to the global genome nucleotide excision repair (GG-NER) pathway, where it acts as a primary sensor of UV-induced DNA lesions. Loss-of-function mutations in the *XPC* gene lead to a photosensitive phenotype, with marked accumulation of unrepaired DNA damage and a dramatically elevated risk (10,000-fold) of skin cancer. However, understanding the molecular signaling mechanisms associated with XP-C has been hindered by the lack of reproducible disease models. Here, we overcome this challenge using our genetically engineered human XPC knockout (KO) keratinocytes, the predominant cell type affected by UV radiation. To uncover upstream signaling changes associated with the absence of XPC expression, we quantified protein tyrosine kinase (PTK) activity one-hour post-UVB exposure. XPC KO keratinocytes showed significant dysregulation of PTK activity on ∼100 phosphosites compared to controls. Complementary mass spectrometry (MS)-based quantitative proteomic analysis performed 24 hours post-UVB exposure identified a downstream signature comprising 791 differentially expressed proteins in XPC KO cells irradiated compared to non-irradiated counterparts. An integrative bioinformatic assessment of the kinase activity and proteomic data revealed a significant perturbation in type I interferon signaling via the JAK/STAT pathway in XPC-deficient keratinocytes, which is further exacerbated by UVB exposure. These findings were validated by western blot analysis, establishing a novel disease-associated molecular signature. Given the central role played by JAK/STAT signaling in inflammatory processes, our results implicate this pathway as a key mediator of XP-C’s hypersensitivity, thereby highlighting its potential as a therapeutic target to alleviate the disease pathology.

## Introduction

Prolonged exposure to solar UV radiation primarily leads to skin aging and an increases risk of developing cancer^1^. This is particularly evident in individuals with xeroderma pigmentosum (XP), a genetic disorder characterized by a deficiency in the nucleotide excision repair (NER) pathway, which is a crucial system for correcting UV-induced DNA damage^2^. The process of NER involves a group of more than 40 proteins that work in cascading manner to repair UV-induced DNA lesions, such as 6–4 pyrimidine pyrimidone photoproducts and cyclobutane pyrimidine dimers (CPDs). This process can split into five main enzymatic stages: (i) DNA damage recognition, (ii) Unwinding of the DNA double helix forming a bubble structure, (iii) Verification of the DNA damage followed by the incision of the single stranded DNA harboring the UV lesion at both 5′ and 3′ ends, (iv) Filling the DNA gap via repair replication, utilizing the intact strand as a template, and ultimately, (v) Sealing the newly synthesized DNA patch through ligation^3^. Perturbation in any of the stages of the NER process can lead to the onset of one of the seven Xeroderma Pigmentosum (XP) complementation groups (XP-A to XP-G)^4^. XPC encodes a key protein involved in initiating the GG-NER. Remarkably, mutations in the *XPC* gene are the most prevalent genetic variation observed in XO. Genodermatosis, also referred to as XP-C disease (OMIM# 278,720)^5^. This protein plays a major role in initiating the GG-NER pathway by identifying and attaching to DNA helical deformities that occur on the undamaged DNA strand opposite to the photoproducts produced by UV radiation. XP-C patients show an increased sensitivity to UV light accompanied by an accumulation of DNA damage. This disorder is often termed as skin cancer prone, as patients have a 2,000 to 10,000 folds risk of developing melanomas and non-melanoma skin cancers (NMSCs) at a young age compared to unaffected individuals^6^. It is noteworthy that till date, crucial to that there is no cure for XP-C syndrome, where the central focus relies on preventative measures using specialized creams against UV exposure^7^. Moreover Moreover, a detailed understanding of the molecular mechanisms and cellular signaling events underlying this pathological condition is still lacking. To get insight into this, our research group has already established reliable human skin cell models for XP-C disease. In fact, we previously generated, using CRISPR-Cas9 system, XP-C disease models using the major three skin cell types (keratinocytes, fibroblasts, and melanocytes), being immortalized from the human skin, and mimicking XP-C disease in terms of major phenotype characteristics: photosensitivity and absence of photoproducts repair induced by UV exposure^8^.

Here, and upon using the established XPC KO keratinocytes model, we aimed at deciphering a precise dysregulated molecular signaling pathway linked to XP-C disease. Using an integrated phosphoproteomics and total proteomics workflow, we profiled early signaling events (1-hour post-UVB exposure) and downstream proteomic changes (24 hours post-exposure). We identified and upregulated activation of JAK/STAT pathway activation and the downstream type I interferon signaling components in XPC KO keratinocytes under both basal conditions and following UVB exposure. These observations suggest that the constitutive activation of inflammatory signaling pathways may underline the pathophysiology of XP-C, beyond its classical role as a DNA repair disorder.

## Results

### 1. Loss of XPC induces a UVB hypersensitive phenotype Over Time in Human Keratinocytes Aberrant phosphorylation of the JAK/STAT Signaling components in XPC Knockout Keratinocytes Following 1 Hour of UVB irradiation

Since XPC loss induces a phenotype hypersensitive to UVB in keratinocytes^8,9^, we, therefore, aimed at characterizing the molecular events underlying such phenotype. In a first step, we compared the viability of XPC KO cells versus wild-type (control) cells following exposure to UVB irradiation at a dose of 100 J/m^2^ over a time period of 48 hours (Figure 1). After 24 hours, we observed a 20% reduction in XPC KO cells viability compared to control cells. Intriguingly, this difference increased to 50% after 48 hours. This could be attributed to the inability of XPC KO cells to initiate an effective DNA repair process, unlike the control cells which showed an improved viability at 48 hours.

**Figure 1.**
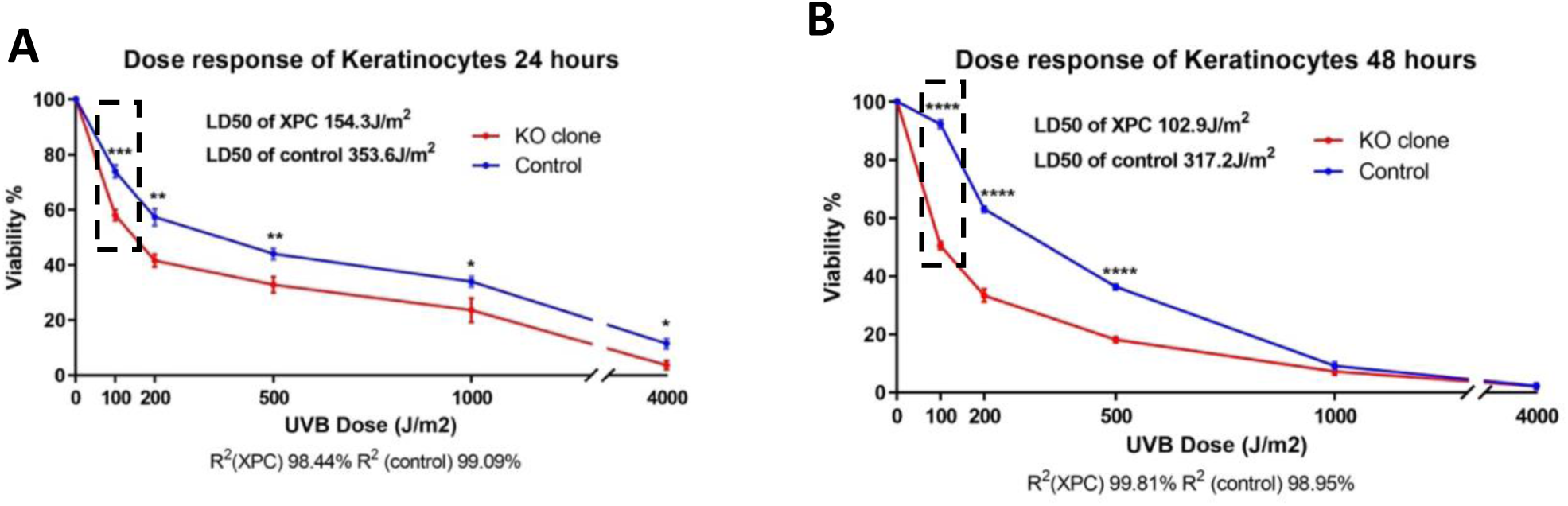
Loss of XPC increases UVB sensitivity over time in N/TERT-2G keratinocytes. Dose–response curves show cell viability following UVB irradiation at the indicated doses, measured at 24 h (left) and 48 h (right). XPC knockout (KO) cells exhibit greater sensitivity than control cells. The 100 J/m² dose is highlighted: at 24 h, the viability difference between KO and control cells is around 20%, but by 48 hours, this gap widens further to 50%. The 100 J/m2 dose (used as the comparative reference for subsequent proteomic analyses). Data are mean ± s.e.m. of three independent experiments (N = 3). *p < 0.05; **p < 0.01; ***p < 0.001; ****p < 0.0001 (unpaired t-test). Figure adapted from Nasrallah et al., Scientific Reports (2024), doi: 10.1038/s41598-024-81675-6, licensed under CC BY 4.0.

### 2. Aberrant phosphorylation of the JAK/STAT Signaling components in XPC Knockout Keratinocytes Following 1 Hour of UVB irradiation

Observing that XPC KO cells exhibit increasing hypersensitivity to UVB irradiation over time prompted us to elucidate the molecular and signaling events underlying this phenomenon. Our workflow strategy integrated two approaches (Figure 2). In the first we used phosphoproteomics analysis to identify activated phosphorylation sites and predict potential kinases while the second we employed total proteomics analysis to unravel deregulated signaling cascades.

**Figure 2.**
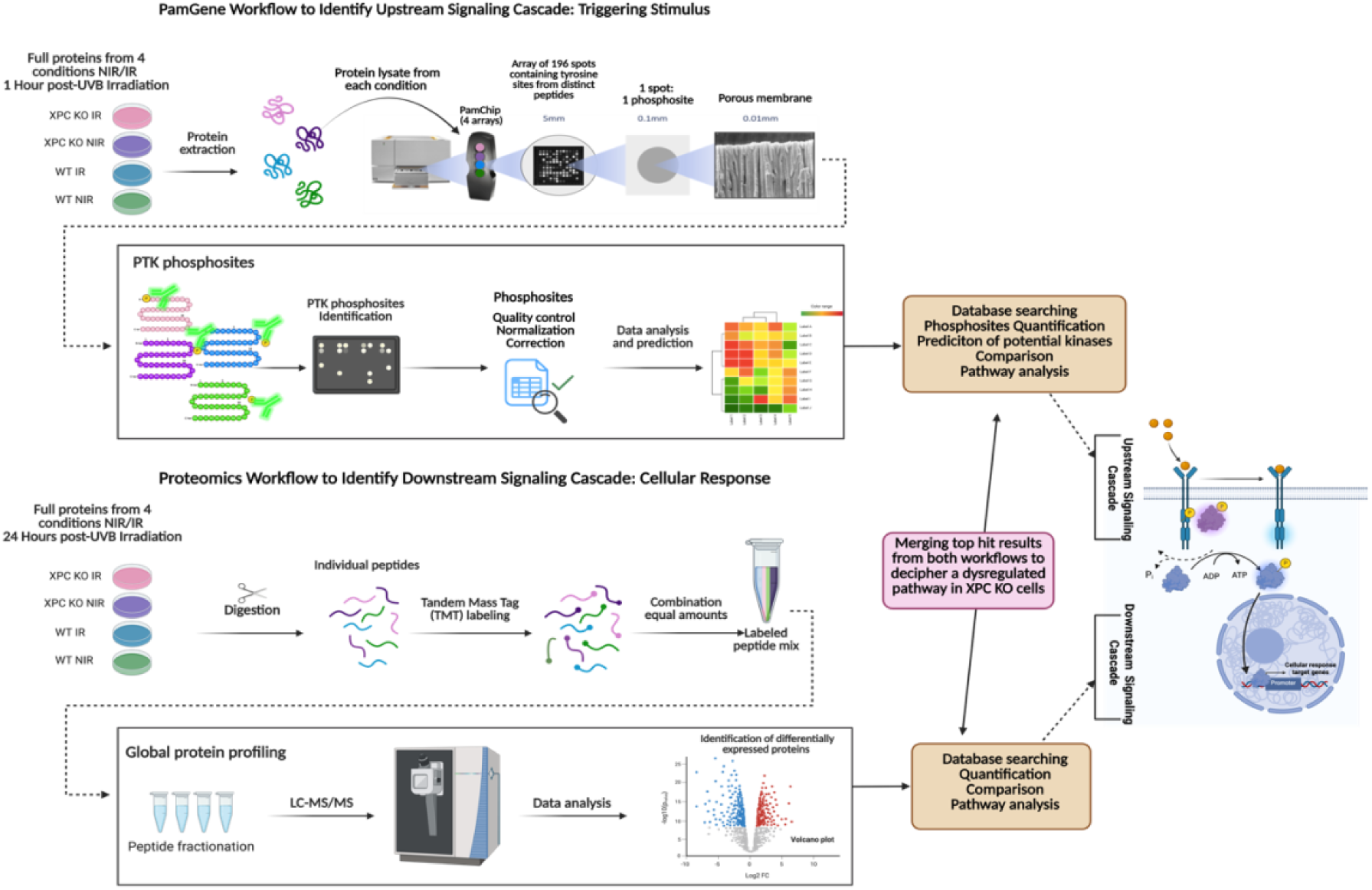
A Novel Strategy for Identifying Dysregulated Signaling Pathways in XPC Knockout Cells Upon UVB Irradiation. A combined phosphoproteomics and global protein profiling approach enables the identification of dysregulated signaling pathways in XPC knockout (KO) cells following UVB irradiation. (Top) Phosphotyrosine kinase (PTK) phosphosite identification workflow, including protein extraction, phosphotyrosine enrichment, array-based detection, and data analysis for kinase prediction and pathway mapping. (Bottom) Global protein profiling workflow involving protein digestion, tandem mass tag (TMT) labeling, peptide fractionation, LC-MS/MS analysis, and identification of differentially expressed proteins. Integrating key findings from both workflows facilitates a systems-level understanding of disrupted signaling cascades in XPC KO cells.

Pamgene kinase activity profiling was applied to evaluate the overall activity of protein tyrosine kinases (PTKs). Kinomic profiles were assessed in XPC KO keratinocytes versus wild-type control cells at basal state as well as 60 mins following UVB irradiation respecting the same experimental conditions applied in Figure 1. Following the quantification of the signals for the four conditions, it was crucial to decipher and quantify their dysregulated phosphosites. Following identification, quantification, and quality control filtering, this procedure unraveled 195 identified PTK phosphosites (excluding background system effects) we then generated (Figure 3)

**Figure 3.**
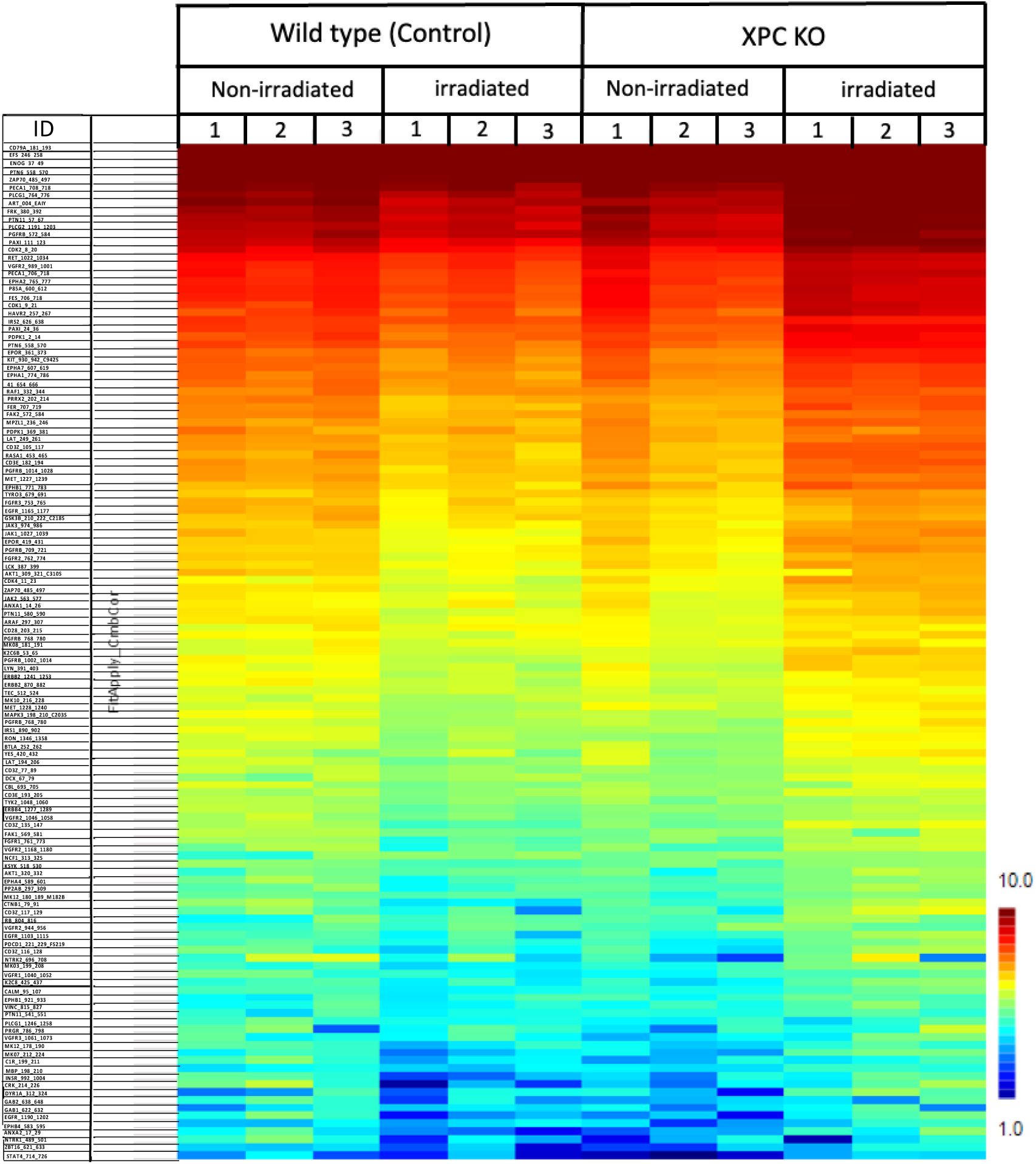
Differential Phosphorylation Signatures in Wild-Type (control) and XPC KO N/TERT-2G keratinocyte cells following 1 hour after UVB Irradiation. Heatmap showing differential phosphorylation profiles across wild-type (control) and XPC knockout (KO) cell lines under non-irradiated and irradiated conditions. Phosphopeptide intensities are represented as a gradient from low (blue) to high (red), highlighting condition-specific and genotype-specific signaling responses. Biological triplicates (N=3) are shown for each group. Key phosphosites are annotated on the y-axis, and samples are grouped along the x-axis according to treatment and genotype.

The phosphorylation profiles varied significantly between the XPC KO cells and wild type (control) cells under the experimental conditions examined (Figure 4A and 4B). For instance, 64 PTK phosphosites were downregulated in UVB irradiated versus non irradiated wild type control cells (WT IR vs WT NIR). On the other hand, 94 PTK phosphosites were upregulated in UVB irradiated versus non irradiated XPC KO cells (XPC KO IR vs XPC KO NIR). Interestingly, 104 PTK phosphosites were upregulated in UVB-irradiated XPC KO cells versus UVB-irradiated wild-type cells (XPC KO IR vs WT IR). These observations suggested a potential role for XPC in regulating the phosphorylation status of PTKs.

**Figure 4.**
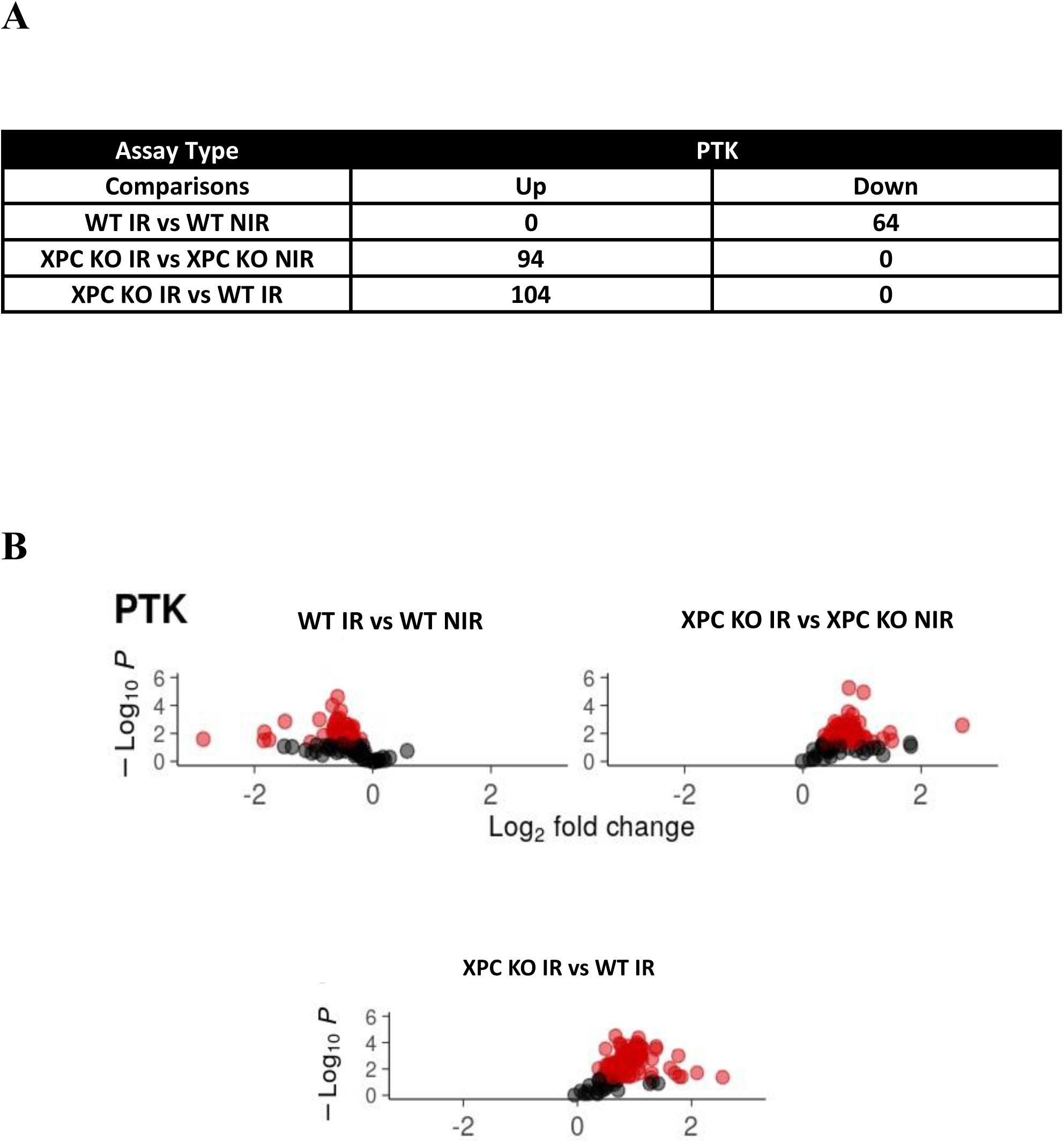
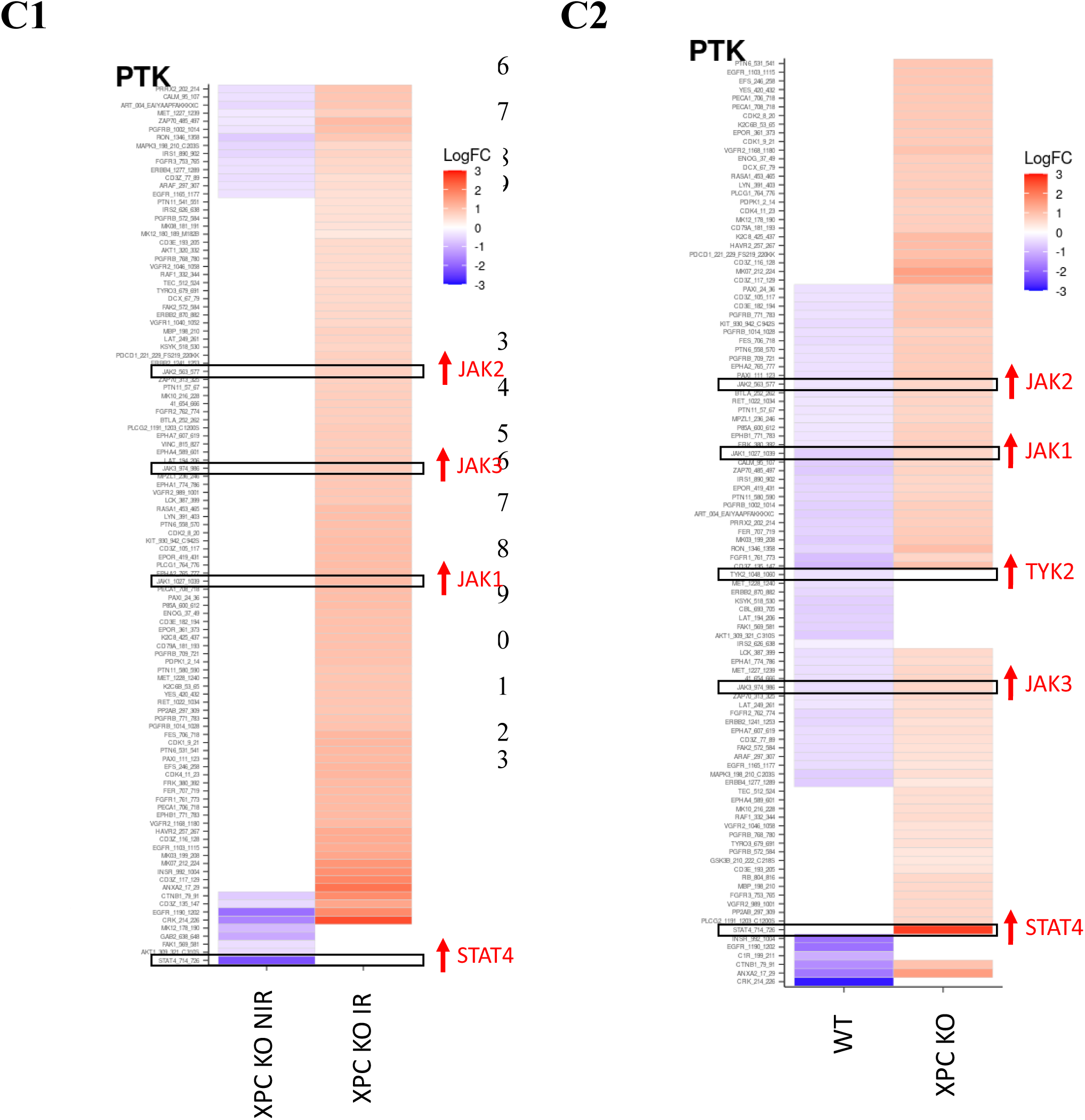
Alterations in PTK Phosphorylation Dynamics Following 1 hour of UVB Irradiation in Wild-Type (control) and XPC KO N/TERT-2G Cells. A). Table showing the number of significantly altered PTK phosphosites across various conditions (WT IR vs WT NIR, XPC KO IR vs XPC KO NIR, XPC KO IR vs WT IR). B) Volcano plots illustrating the distribution of differentially phosphorylated PTK phosphosites in response to UVB irradiation in various conditions. Comparisons shown: WT IR vs WT NIR, XPC KO IR vs XPC KO NIR, and XPC KO IR vs WT IR. Red points indicate significantly altered phosphorylation (p < 0.05). C1). Heatmap presentation of differentially altered PTK peptide phosphosites in wild-type versus XPC KO keratinocytes following UVB irradiation. This heatmap compares both wild-type UVB irradiated versus non-UVB irradiated keratinocytes (noted as WT) along with XPC KO UVB irradiated versus non-UVB irradiated keratinocytes (noted as XPC KO) for PTK peptide phosphosites. JAK/STAT family: JAK1, JAK2, TYK2, JAK3, and STAT4 (in red) were further selected for future validation experiments. *** p< 0.001, unpaired t-test. The results presented are the mean of three biological replicates (N=3). C2). Heatmap presentation of differentially altered PTK peptide phosphosites in XPC KO keratinocytes at basal state versus UVB irradiation. This heatmap compares XPC KO non-UVB irradiated versus wild type non-UVB irradiated keratinocytes (noted as XPC KO NIR) along with XPC KO UVB irradiated normalized to wild type UVB irradiated keratinocytes (noted as XPC KO IR). JAK/STAT family: JAK1, JAK2, JAK3, and STAT4 (in red) were further selected for future validation experiments. The logFC ranged from +3 to −3 for significant peptide phosphorylation sites for PTK. *** p< 0.001, unpaired t-test. The results presented are the mean of three biological replicates (N=3).

In a further step, we designed a heatmap comparing phosphosites in IR/NIR wild-type control cells versus IR/NIR XPC KO cells (Figure 4C1) as well as another heatmap comparing phosphosites in NIR XPC KO /NIR wild-type cells versus IR XPC KO /IR wild-type cells (Figure 4C2). Figure 4C1 revealed a significant upregulation in the phosphorylation status across a broad range of PTK peptide phosphosites following XPC knockout. Among the identified phosphosites, were those of the JAK/STAT family: JAK1, JAK2, TYK2, JAK3, as well as STAT4. Interestingly, the phosphorylation status of those members of the JAK/STAT signaling pathway were also upregulated in response to UVB irradiation as monitored in IR-versus NIR-XPC KO cells (Figure 4C2). These observations highlight a potential role for the deregulated JAK/STAT signaling pathway in mediating the XPC KO phenotype in response to UVB irradiation.

The upregulation of these players reflects the robustness of the dysregulation of this pathway and can exclude the false positive effect. Thus, JAK/STAT signaling pathway seemed to be a target of dysregulation for XPC KO keratinocytes following UVB irradiation. The phosphosite analysis showed the dysregulation of JAK/STAT family members in XPC KO keratinocytes post-UVB irradiation.

### 2. Global Proteomic Profiling Reveals Activation of Type I Interferon Downstream Effectors 24 Hours After UVB Irradiation in XPC-Deficient Keratinocytes

Following the second approach of our workflow strategy, the total proteomic signature was assessed in both XPC KO keratinocytes versus wild-type cells following the same experimental conditions applied in Figure 1. Protein samples derived from non-irradiated wild-type cells, UVB-irradiated wild-type cells, non-irradiated XPC KO cells and UVB-irradiated XPC KO cells were prepared then analyzed by mass spectrometry as mentioned in the materials and methods section.

A total of 6699 proteins were identified to be deregulated. For each differential analysis, identical thresholds: log2(FoldChange) = 0.3 and p-value = 0.01 were selected. This corresponds to an enrichment of at least 1.23 times in one condition compared to the other. Volcano plots for each differential comparison were further constructed using log2 foldchange (log2FC) on the x-axis and -log10pvalue on the y-axis for better scale fitting and to visualize the distribution of differentially expressed proteins between conditions. Statistical analyses were then carried out to identify the proteins that were differentially abundant between the conditions compared in pairs. 390 proteins were identified to be differentially abundant in wild-type keratinocytes following UVB irradiation compared to non-irradiated ones (Figure 5A). When comparing XPC KO UVB irradiated and non-UVB irradiated keratinocytes (XPC KO IR vs XPC KO NIR), 791 proteins were identified to be significantly dysregulated (upregulated or downregulated), representing more than 10% of the quantified proteins (Figure 5B). When comparing XPC KO UVB irradiated vs wild type UVB irradiated keratinocytes (XPC KO IR vs WT IR), 413 proteins were found to be differentially abundant (upregulated or downregulated) (Figure 5C). The highly upregulated proteins (log2foldchange) were set on the top list whereas the most downregulated proteins were set on the bottom list. It is noteworthy that IR XPC KO normalized to IR-wild-type was used as a reference. The top list proteins included Interferon-induced GTP-binding protein (MX1), Serpin Family B Member 3 (SERPINB3), S100 calcium-binding protein A12 (S100A12), Interferon Induced Protein With Tetratricopeptide Repeats 2 (IFIT-2), Interferon Induced Protein With Tetratricopeptide Repeats 3 (IFIT-3), Interferon Induced Protein With Tetratricopeptide Repeats 1 (IFIT-1), Interferon-induced GTP-binding protein (MX2), Suprabasin (SBSN), Interleukin 1 beta (IL-1β), S100 calcium-binding protein A7 (S100A7), Calmodulin Like 5 (CALML5), 2’-5’-Oligoadenylate Synthetase 1 (OAS1), Interferon Regulatory Factor 9 (IRF9), and Interferon-stimulated gene 15 (ISG15) (Figure 6). STRING protein-protein interaction network software (Figure 7A) was implemented for the selected top cluster of dysregulated proteins along with JAK/STAT family, including JAK1, JAK2, JAK3, STAT1, STAT2, and STAT3, these proteins were also identified to be deregulated using the proteomics analysis but were not on the top list to check whether these proteins form an interaction network. Interestingly, the results showed a robust protein-protein interaction network between this cluster of dysregulated proteins and the JAK/STAT family members, except for suprabasin, which was not linked to the network. To better predict the signaling pathway in which this protein network is involved, g:Profiler software was applied (Figure 7B).

**Figure 5.**
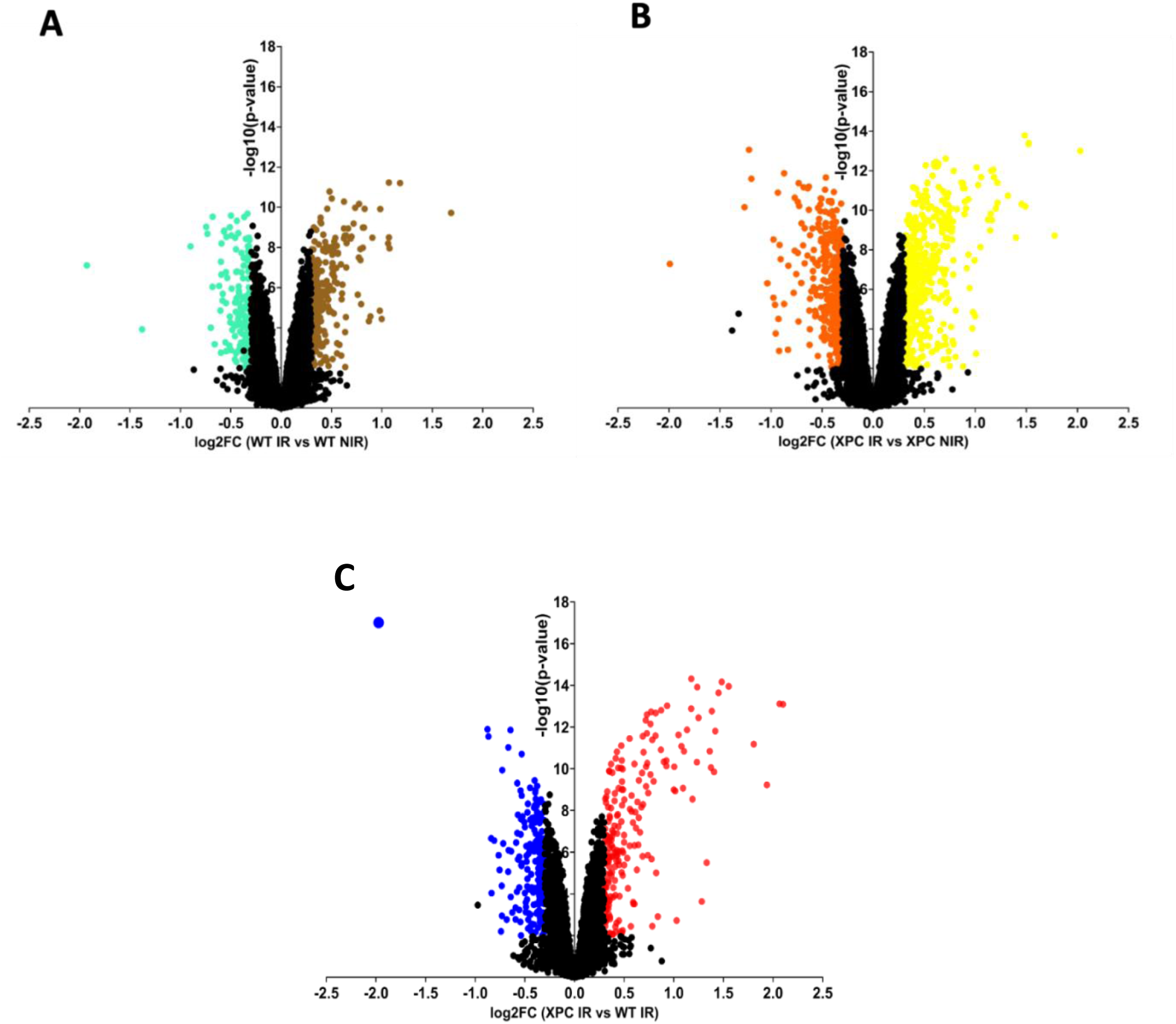
Volcano plot representation of differentially expressed proteins across keratinocyte genotypes and UVB exposure conditions. Volcano plots illustrate the significantly upregulated and downregulated proteins identified in four pairwise comparisons involving N/TERT-2G wild-type and XPC knockout (KO) keratinocytes, under basal conditions and following UVB irradiation. (A) Wild-type keratinocytes: UVB-irradiated versus non-irradiated (WT IR vs WT NIR). (B) XPC KO keratinocytes: UVB-irradiated versus non-irradiated (XPC IR vs XPC NIR). (C) UVB-irradiated XPC KO versus UVB-irradiated wild-type keratinocytes (XPC IR vs WT IR). Statistical thresholds applied across all comparisons were log₂ (fold change) ≥ ±0.3 (corresponding to ≥1.23-fold change) and p-value ≤ 0.01. False discovery rates (FDR) were controlled at approximately 1% for each comparison: WT UVB vs. non-UVB (1.14%), XPC KO UVB vs. non-UVB (1.08%), XPC KO vs. WT at baseline (1.34%), and XPC KO vs. WT post-UVB (1.18%). Volcano plots were generated using GraphPad Prism v8, with log₂ (fold change) on the x-axis and –log₁₀(p-value) on the y-axis to visualize the distribution and significance of differentially expressed proteins. Significance was assessed using both unpaired and paired t-tests; ***p < 0.001. Data represent the mean of four biological replicates (N = 4) per condition.

**Figure 6.**
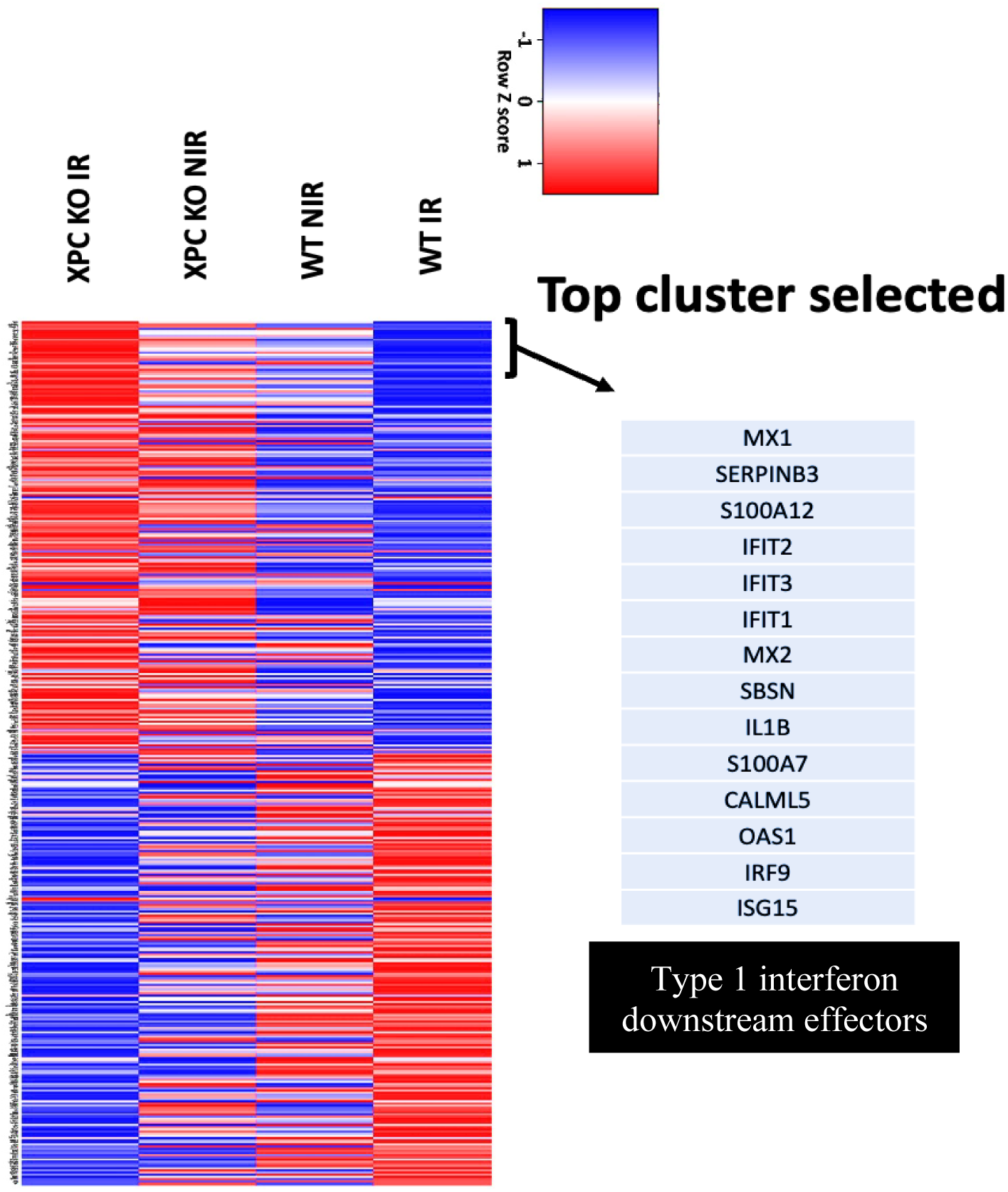
Heat map presentation of the significantly and differentially upregulated and downregulated proteins in four various conditions encompassing N/TERT-2G wild-type and XPC KO keratinocytes at basal state and following UVB irradiation. This heatmap presentation shows the differential expression profile of 413 proteins (arranged from the highly upregulated on the toplist to the downregulated on the bottom list according to the condition XPC KO IR vs WT IR used as a reference). Each row is a protein and each column corresponds to a specific condition starting from XPC KO UVB irradiated vs wild-type UVB irradiated (XPC KO IR vs WT IR), XPC KO UVB irradiated vs non-UVB irradiated (XPC KO IR vs XPC KO NIR), XPC KO non-UVB irradiated vs wild-type non-UVB irradiated (XPC KO NIR vs WT NIR), and wild-type UVB irradiated vs non-UVB irradiated (WT IR vs WT NIR). The 14 top list proteins selected according to the condition XPC KO IR vs WT IR compared to the other three conditions were Interferon-induced GTP-binding protein (MX1), Serpin Family B Member 3 (SERPINB3), S100 calcium-binding protein A12 (S100A12), Interferon Induced Protein With Tetratricopeptide Repeats 2 (IFIT-2), Interferon Induced Protein With Tetratricopeptide Repeats 3 (IFIT-3), Interferon Induced Protein With Tetratricopeptide Repeats 1 (IFIT-1), Interferon-induced GTP-binding protein (MX2), Suprabasin (SBSN), Interleukin 1 beta (IL-1β), S100 calcium-binding protein A7 (S100A7), Calmodulin Like 5 (CALML5), 2’-5’-Oligoadenylate Synthetase 1 (OAS1), Interferon Regulatory Factor 9 (IRF9), and Interferon-stimulated gene 15 (ISG15). The FunRich Version 3.1.4 was used to construct this heatmap. The Row Z score fluctuated from −1 to 1. *** p< 0.001, unpaired and paired t-test. The results presented are the mean of four biological replicates (N=4).

**Figure 7.**
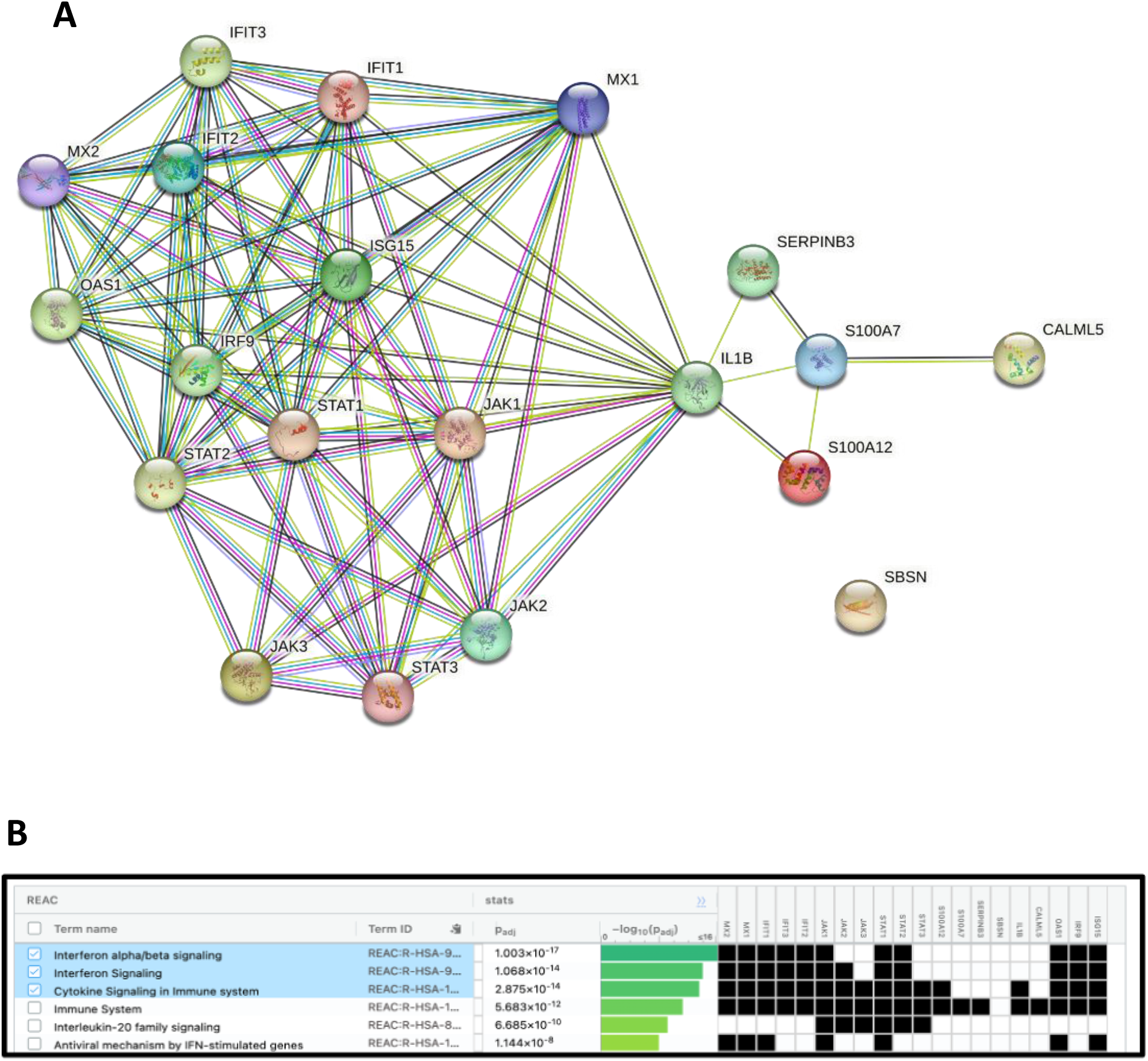
Integrated Protein-Protein Interaction Network and Enriched Biological Pathways of Dysregulated Proteins in XPC KO UVB-Irradiated Cells Compared to Wild-Type UVB-Irradiated Cells, Highlighting JAK/STAT Signaling. (A). This figure presents an integrated view of the top dysregulated proteins (upregulated) in XPC knockout (KO) cells following UVB irradiation compared to wild-type (WT) UVB-irradiated cells. The protein-protein interaction (PPI) network, constructed using STRING database v11.5, reveals extensive associations among 13 key upregulated proteins—MX1, MX2, SERPINB3, S100A12, IFIT1, IFIT2, IFIT3, IL-1β, S100A7, CALML5, OAS1, IRF9, and ISG15—along with JAK/STAT family members (JAK1, JAK2, JAK3, STAT1, STAT2, STAT3), previously identified as a dysregulated upstream kinase cluster. The network underscores the central role of immune and interferon-related signaling. (B). In parallel, enrichment analysis using g:Profiler v2.1 revealed significant clustering of these proteins into common biological pathways based on Reactome database annotations. The most prominent pathways include Type I interferon-alpha/beta signaling, general interferon signaling, and cytokine signaling in the immune system. Together, these findings suggest a strong immune and interferon-driven response in XPC KO cells upon UVB exposure, mediated in part by JAK/STAT signaling.

Our results unraveled Type I Interferon alpha/beta signaling pathway to be on the top list including MX1, MX2, IFIT1, IFIT2, IFIT3, JAK1, STAT1, STAT2, OAS1, IRF9, and ISG15. The second top hit was Interferon signaling including MX1, MX2, IFIT1, IFIT2, IFIT3, JAK1, JAK2, STAT1, STAT2, OAS1, IRF9, and ISG15. The third top hit was Cytokine Signaling in Immune system including MX1, MX2, IFIT1, IFIT2, IFIT3, JAK1, JAK2, JAK3, STAT1, STAT2, STAT3, S100A12, IL1β, OAS1, IRF9, and ISG15.

### 3. Validation of Upstream and Downstream Components of Type I Interferon Signaling Through the JAK/STAT Pathway in XPC Knockout Keratinocytes

The above observations indicated an upregulated role for JAK/STAT signaling pathway in UVB-irradiated XPC KO keratinocytes. Accordingly, a set of experiments were conducted to validate this observation.

In a first step, we assessed the phosphorylation/expression status of several STAT proteins including STAT1, STAT2 and STAT3. Western blot analysis was therefore performed using antibodies against alpha phosphoTyr-701-STAT-1 (91 kDa), phosphoTyr-690-STAT-2 (97 kDa), and phosphoTyr-705-STAT-3 (86 kDa) (Figure 8A). In the case of alpha phosphoTyr-701-STAT-1 (91 kDa), the expression/phosphorylation level was strikingly higher in UVB-irradiated XPC KO cells versus UVB-irradiated wild-type control cells (Figure 8B). Moreover, the expression status was significantly higher in UVB-irradiated XPC KO cells versus non-irradiated XPC KO cells (Figure 8B). Similarly in the case of phosphoTyr-690-STAT-2 (97 kDa), UVB-irradiated XPC KO cells showed the highest expression/phosphorylation level. This was 5x higher than that of UVB-irradiated wild-type cells, 2.5x more than that exhibited by non-irradiated wild-type cells, and 1.11x greater than that of non-irradiated XPC KO cells (Figure 8C). Further, in the case of phosphoTyr-705-STAT-3 (86 kDa), the expression/phosphorylation levels were substantially higher in XPC KO cells versus wild-type control cells in absence or presence of UVB irradiation (Figure 8D).

**Figure 8.**
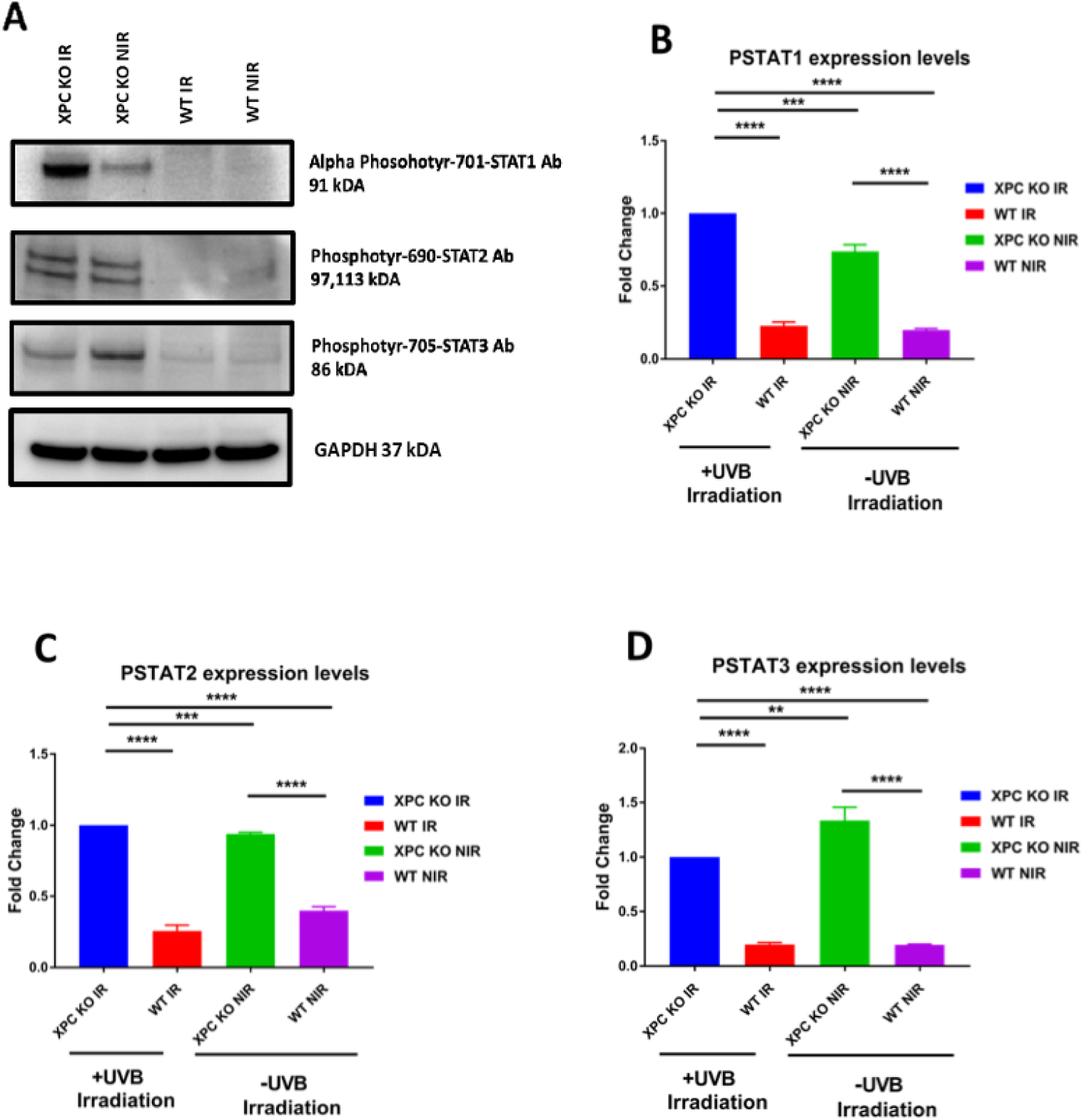
Increased phosphorylation of STAT proteins in XPC knockout keratinocytes under basal conditions and after UVB exposure. (A) Western blot analysis showing increased phosphorylation of STAT family members specifically phospho-Tyr701-STAT1 (91 kDa), phospho-Tyr690-STAT2 (97 kDa), and phospho-Tyr705-STAT3 (86 kDa) in XPC-deficient keratinocytes compared to wild-type cells, both at baseline and post-UVB irradiation. (B–D) Quantification of fold changes in expression levels for (B) phospho-Tyr701-STAT1, (C) phospho-Tyr690-STAT2, and (D) phospho-Tyr705-STAT3. Protein lysates were resolved via SDS-PAGE and transferred to nitrocellulose membranes. Detection was performed using phospho-specific antibodies, with GAPDH (37 kDa) serving as a loading control. Fold change values were normalized to GAPDH. Statistical significance was assessed using unpaired and paired t-tests (***p < 0.001). Data represent mean values from three independent replicates (N = 3).

In a second step, western blot analysis was used to assess the expression levels of several downstream effectors (MX1, MX2, IFIT1, IFIT3, IRF9, and ISG15) of the JAK/STAT signaling pathway (Figure 9). Interestingly, the expression levels of IRF-9 (Figure 9B), IFIT1 (Figure 9C), IFIT3 (Figure 9D), MX1 (Figure 9F), ISG15 (Figure 9H) and MX2 (Figure 9I) were strikingly higher in XPC KO cells versus wild-type cells either in absence or presence of UVB irradiation. Intriguingly, the expression levels of those genes were significantly upregulated following exposure of XPC KO cells to UVB-irradiation (Figure 9). For IRF-9 (Figures 9A and 9B), the results showed a significant upregulation of IRF-9 in XPC KO UVB irradiated keratinocytes (5.5 folds), in XPC KO non-UVB irradiated keratinocytes (3.3 folds), and a slight increase for wild-type UVB irradiated keratinocytes (1.5 folds) compared to the reference utilized which was wild-type non-UVB irradiated keratinocytes set at a constant 1-fold to compare. For IFIT1 (Figure 9C and 9E), the results showed a significant upregulation of IFIT1 in XPC KO UVB irradiated keratinocytes (15.5 folds), in XPC KO non-UVB irradiated keratinocytes (7.5 folds), and a constant expression level for wild-type UVB irradiated keratinocytes (1-fold) compared to the reference utilized which was wild-type non-UVB irradiated keratinocytes set at a constant 1 fold to compare. For IFIT3 (Figure 9D and 9E), the results showed a significant upregulation of IFIT3 in XPC KO UVB irradiated keratinocytes (17.5 folds), in XPC KO non-UVB irradiated keratinocytes (5 folds), and a slight increase for wild-type UVB irradiated keratinocytes (1.75 folds) compared to the reference utilized which was wild-type non-UVB irradiated keratinocytes set at a constant 1-fold to compare. For MX1 (Figure 9F and 9E), the results showed a significant upregulation of MX1 in XPC KO UVB irradiated keratinocytes (15.7 folds), in XPC KO non-UVB irradiated keratinocytes (5.5 folds), and a constant expression level for wild-type UVB irradiated keratinocytes (1-fold) compared to the reference utilized which was wild-type non-UVB irradiated keratinocytes set at a constant 1 fold to compare. For MX2 (Figure 9G and 9I), the results showed a significant upregulation of MX2 in XPC KO UVB irradiated keratinocytes (20.5 folds), in XPC KO non-UVB irradiated keratinocytes (9 folds), and a slight downregulation for wild type UVB irradiated keratinocytes (0.75 folds) compared to the reference utilized which was wild type non-UVB irradiated keratinocytes set at a constant 1-fold to compare. For ISG15 (15 kDa) (Figure 9H and 9G), the results showed a significant upregulation of ISG15 in XPC KO UVB irradiated keratinocytes (19 folds), in XPC KO non-UVB irradiated keratinocytes (2 folds), and a constant expression level for wild-type UVB irradiated keratinocytes (1-fold) compared to the reference utilized which was wild-type non-UVB irradiated keratinocytes set at a constant 1-fold to compare.

**Figure 9.**
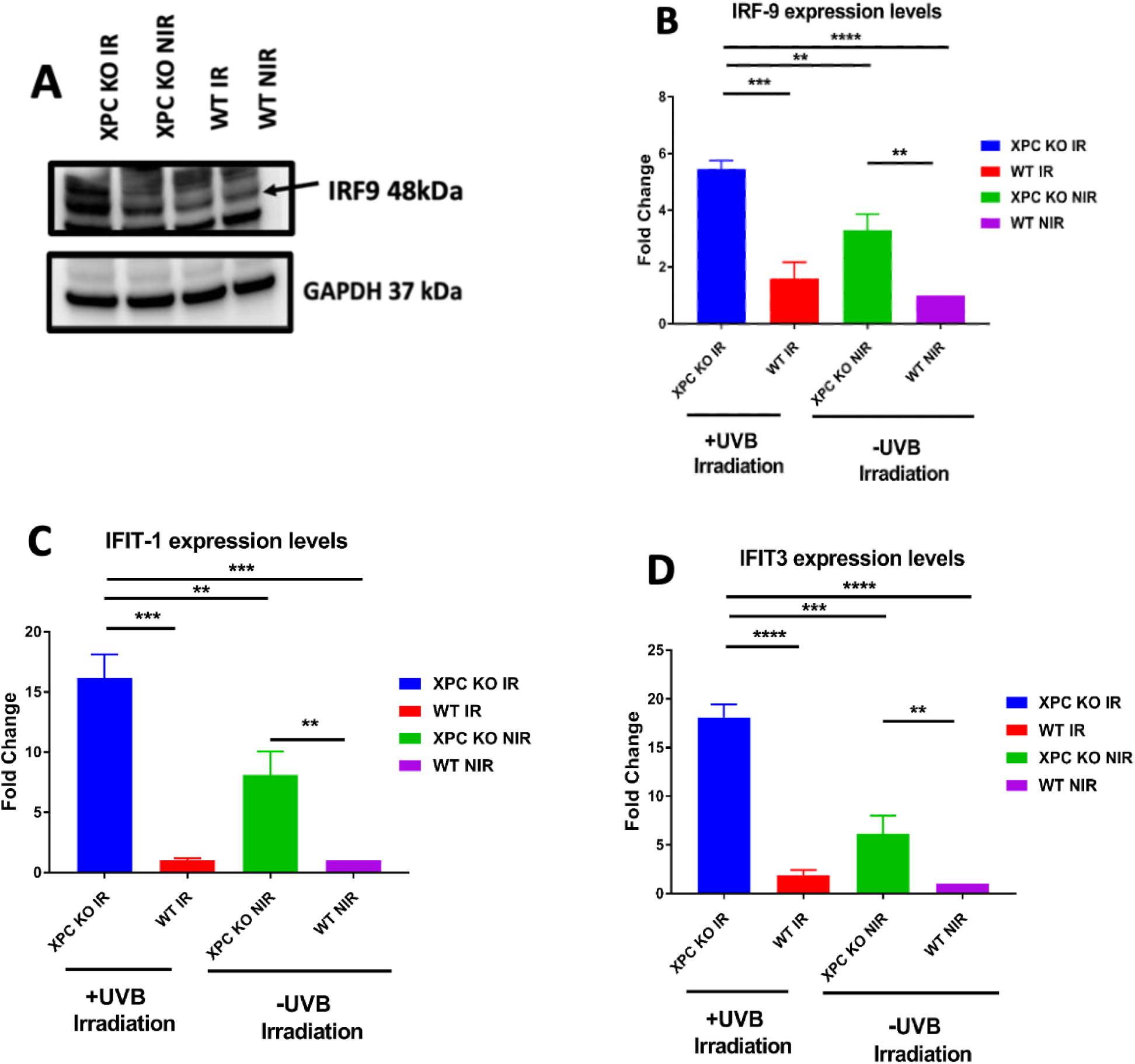
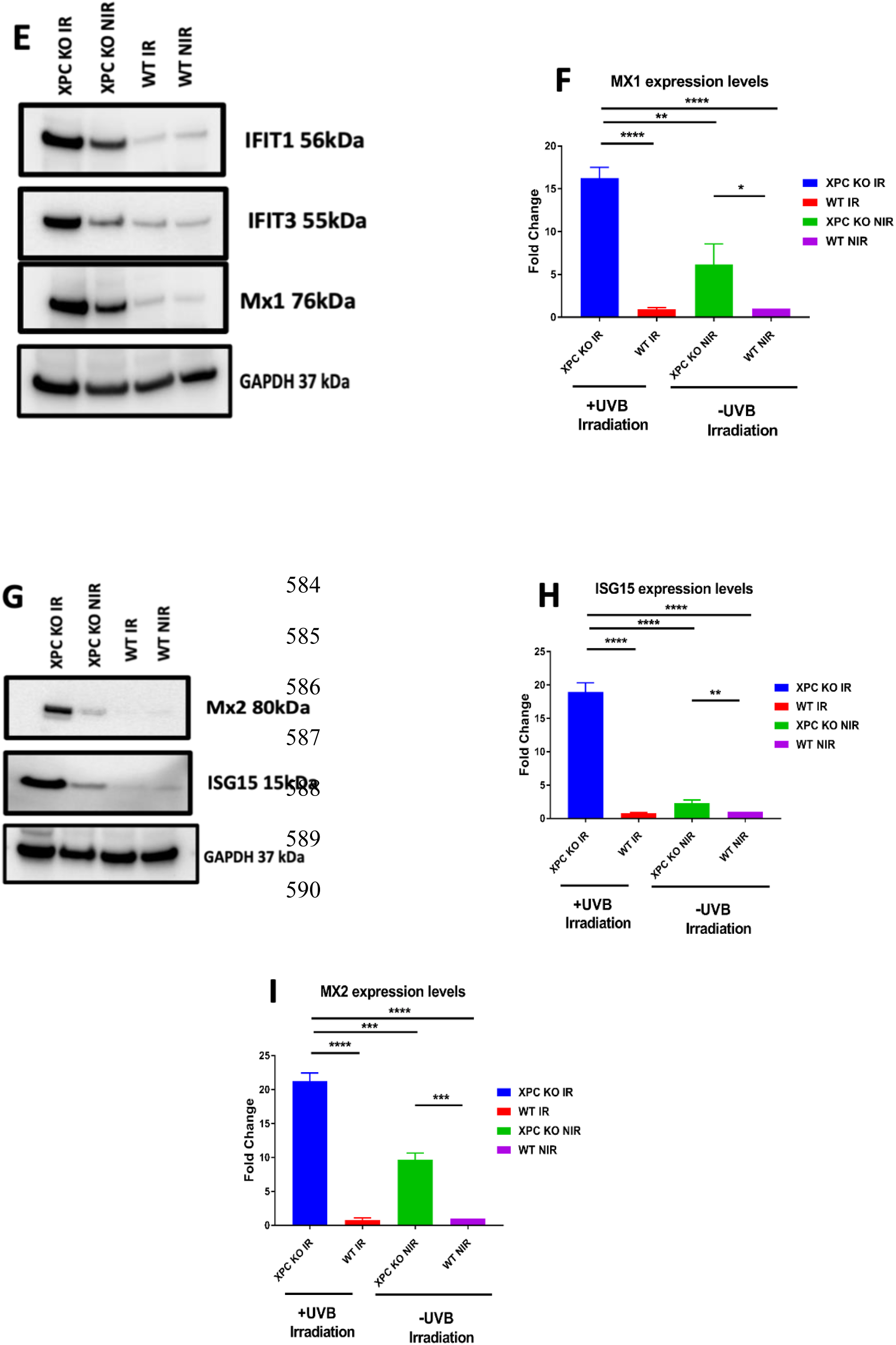
UVB irradiation induces robust activation of Type I interferon downstream effectors in XPC-deficient keratinocytes. Immunoblotting and quantitative analysis reveal elevated expression of Type I interferon downstream effectors in XPC knockout (KO) keratinocytes compared to wild-type (WT) controls following UVB irradiation. (A, E, G) Immunoblots showing expression of IRF9 (48 kDa), IFIT1 (56 kDa), IFIT3 (55 kDa), MX1 (76 kDa), MX2 (80 kDa), and ISG15 (15 kDa) across four experimental groups: XPC KO irradiated (IR), XPC KO non-irradiated (NIR), WT IR, and WT NIR. GAPDH (37 kDa) served as loading control. (B–D, F, H–I) Quantification of fold-change in protein expression for IRF9 (B), IFIT1 (C), IFIT3 (D), MX1 (F), ISG15 (H), and MX2 (I) reveals significantly higher induction in XPC KO cells 24 hours post-UVB exposure compared to other groups. Data are presented as mean ± SEM; statistical analysis performed using paired and unpaired T-test (*P < 0.05; **P < 0.01; ***P < 0.001; ***P < 0.0001). These findings support activation of the JAK/STAT-mediated Type I interferon signaling axis in the absence of functional XPC in response to UVB-induced DNA damage.

Altogether, these observations are in support of a JAK/STAT mediated-upregulation of Type I interferon alpha/beta signaling in XPC KO keratinocytes at basal state as well as following UVB irradiation.

## Discussion

This study aimed at better understanding the molecular mechanisms underlying the phenotypic characteristics of XP-C disease. We succeeded in this report, for the first time, in deciphering a signaling cascade that is dysregulated in XPC KO cells. In fact, we show that the JAK/STAT pathway and its downstream effectors are significantly up regulated in XPC knockout keratinocytes under both basal conditions or following exposure to UVB.

XP-C disease is one of the most prevalent subtypes of xeroderma pigmentosum^10^ and is typically caused by nonsense mutations, frameshifts, or deletions in the *XPC* gene^11^. These mutations often lead to complete loss of the XPC protein and negligible mRNA expression^11^. XP-C cells are profoundly hypersensitive to UVB and exhibit impaired repair of UV-induced DNA lesions such as 6-4 photoproducts (6-4PPs) and cyclobutane pyrimidine dimers (CPDs), due to a defective global genome nucleotide excision repair (GG-NER) mechanism^9^. Given that the epidermis is the primary site of UV exposure, our experimental model was based on keratinocytes: the most abundant epidermal cells and a key target in non-melanoma skin cancers (NMSCs), which occur at rates up to 10,000 times higher in XP-C patients than in the general population.

Despite the clinical importance of keratinocytes in XP-C pathology, the field has lacked reliable experimental models due to the rarity of patient-derived keratinocyte cells and challenges associated with their limited lifespan and absence of genetically isogenic controls^12^. This does not exclude the importance of validating our results on XP-C keratinocytes from patients in the future, once such samples become accessible. Nevertheless, our current strategy provides a valuable platform to facilitate the elucidation of molecular signaling pathways, potentially enabling direct validation and therapeutic development in more advanced models. Additionally, mouse models can offer partial data but are limited by significant interspecies differences in skin biology and difficulties in maintaining primary mouse keratinocytes in vitro as there is no reliable media and protocols to culture them as well as maintain them in culture^13^. To overcome these limitations, we sought to choose the immortalized human N/TERT-2G keratinocyte cell line. This model has a normal karyotype^14^, supports long-term culture^15^, and faithfully recapitulates XP-C phenotypes, including the hypersenisitve phenotype towards UVB irradiation and delayed repair of UVB-induced DNA damage^8^. Its utility for therapeutic research has also been demonstrated. Kobaisi et al. showed that targeting PIK3C3 and LATS1 led to over 20% of repair restoration of 6-4PP DNA damage using this XPC KO model^16^.

To dissect the molecular alterations in XPC-deficient keratinocytes, we implemented a dual-stage proteomics workflow. Phosphoproteomics was used to identify early signaling changes one-hour post-UVB irradiation, while total proteome analysis at 24 hours revealed broader downstream effects. This integrated approach enabled the construction of a high-resolution signaling map, revealing that type I IFN signaling was aberrantly activated in XPC KO cells both at baseline and after UVB exposure through the JAK/STAT axis. This strategy not only illuminates the cellular pathways affected by UVB exposure but also represents a powerful approach for uncovering disease-related mechanisms on a broader scale. By mapping critical signaling disruptions, it has the potential to inform the development of targeted therapies and advance our understanding of complex disease biology.

A major strength of this study was that we combined phosphoproteomics and total proteomics analysis. Phosphoproteomics was used to identify early signaling changes one-hour post-UVB irradiation, while total proteome analysis revealed broader downstream effectors. Despite the fact that this experimental strategy unraveled deregulated expression of many molecular components, a remarkable and significant upregulation of components of JAK/STAT signaling cascade as well as its downstream type I IFN signaling components were upregulated in response to XPC Knockout.

The JAK/STAT pathway (Figure 10) is activated upon cytokine binding to IFNAR1/IFNAR2 receptor complexes, which triggers phosphorylation of TYK2 and JAK1, followed by downstream activation of STAT1, STAT2, and STAT3^17^. Upon UVB exposure, we found a strong induction of pSTAT1 and a modest upregulation of pSTAT2, indicating activation of the type I interferon signaling pathway. Furthermore, pSTAT3 was already elevated under basal conditions in XPC KO keratinocytes, suggesting a constitutive dysregulation. We further confirmed the phosphorylation of specific tyrosine residues: Tyr701 on STAT1α and Tyr690 on STAT2, which are known to be phosphorylated by JAK1 and TYK2, triggering downstream interferon responses^18,19^. Additionally, STAT3 activation involves phosphorylation at Tyr705, and is known to crosstalk with JAK pathway^20^. By quantifying the phosphorylation status of these residues, we confirmed the validity of our hypothesis. These phosphorylated STATs form homodimers or heterodimers and translocate to the nucleus, driving transcription of interferon-stimulated genes (ISGs) via binding to GAS or ISRE elements. In our model, activation of this cascade was accompanied by upregulation of hallmark ISGs such as ISG15, MX1, MX2, IFIT1, and IFIT3, suggesting a persistent inflammatory state in XPC-deficient keratinocytes^17^.

**Figure 10.**
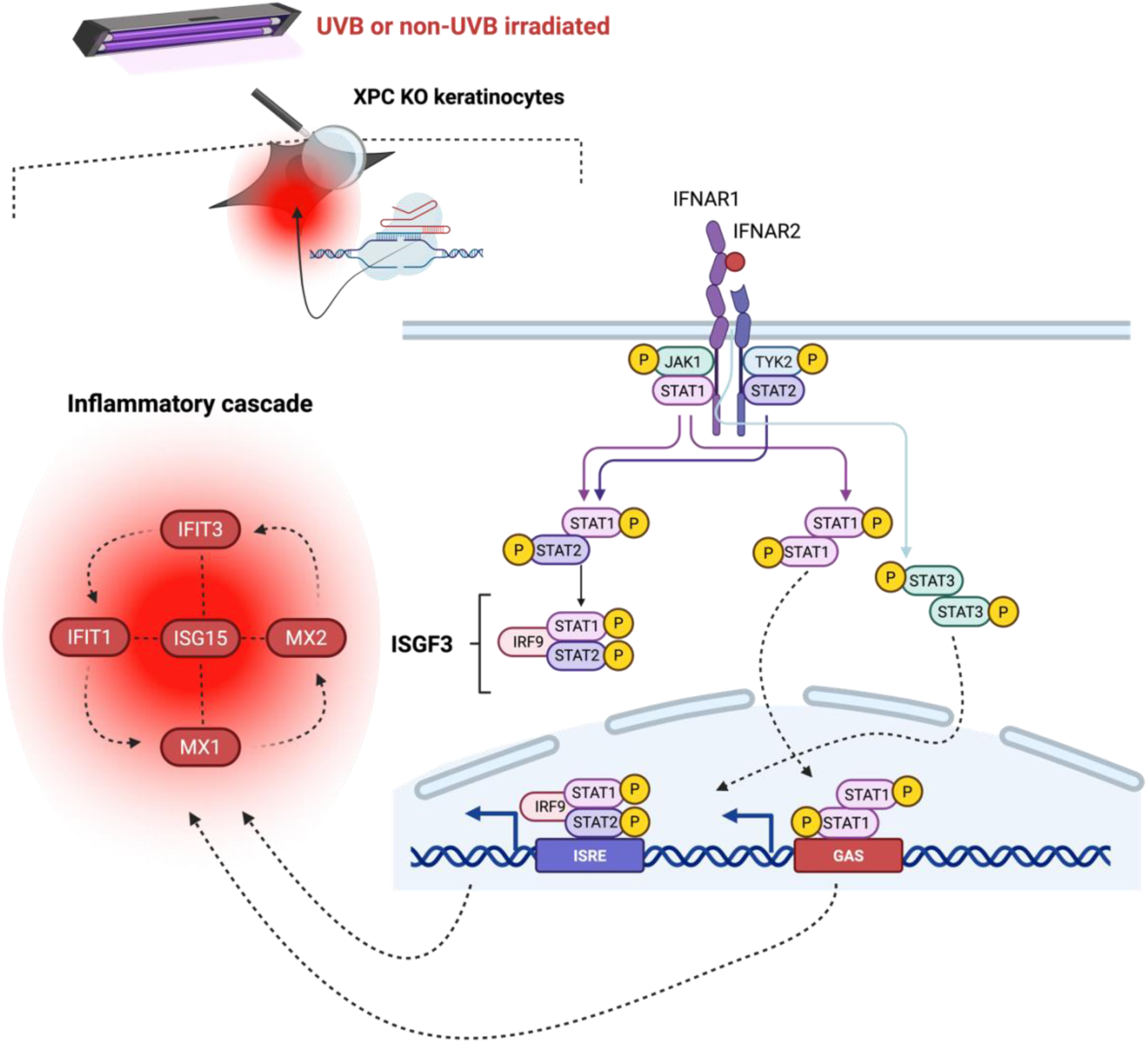
Hypothetical model of Type I interferon pathway activation via JAK/STAT signaling in XPC-deficient keratinocytes. Schematic representation of a proposed mechanism by which Type I IFN signaling is selectively activated in XPC knockout (KO) keratinocytes. Type I interferons signal through the heterodimeric IFNAR1/IFNAR2 receptor complex, which undergoes conformational changes upon ligand binding, initiating phosphorylation of Tyk2 and JAK1. This leads to the activation of STAT1 and STAT2, and occasionally STAT3 via JAK kinases. Activated STATs dimerize to form transcriptionally active complexes. The STAT1–STAT2 heterodimer associates with IRF9 to form ISGF3, which translocates to the nucleus and binds to interferon-stimulated response elements (ISREs), inducing expression of pro-inflammatory genes such as *MX1*, *MX2*, *IFIT1*, *IFIT3*, and *ISG15*. STAT1 and STAT3 homodimers can also independently bind to gamma-activated sequence (GAS) elements to modulate similar inflammatory responses.

While the type I interferon pathway emerged as a central component of the dysregulated signaling landscape in XPC KO keratinocytes, pharmacological inhibition of this pathway remains technically challenging in this context. Preliminary attempts to block key nodes (e.g., JAK1/STAT1) using chemical molecule inhibitors (such as Tofacitinib and Filgotinib) led to non-specific effects due to widespread pathway crosstalk and the extensive perturbation of inflammatory signaling networks following XPC loss and UVB exposure as several other inflammatory pathways including TLR, NF-κB, p-SRC, MAPK, and c-MET have also been shown to be dysregulated through our phosphoproteomics and MS-based quantitative proteomics analyses. Given that hundreds of genes were significantly dysregulated, isolating the contribution of a single pathway without affecting others proved difficult which might explain the incapability to reverse the phenotype of the disease. Future work will benefit from the use of more refined, pathway-specific genetic or pharmacological tools, such as inducible knockdowns or CRISPR interference screens, to dissect causality with higher resolution^21^.

Interestingly, this IFN-driven transcriptional signature mirrors that seen in other UV-related inflammatory skin disorders, including cutaneous lupus erythematosus^22^, psoriasis^23^, and vitiligo^24,25^, and morphea^26,27^, where type I IFN activation contributes to tissue damage and immune dysregulation. These similarities raise the possibility that XP-C could share molecular features with inflammatory skin diseases, extending beyond its classification as a DNA repair disorder. Thus, our findings not only illuminate the contribution of the type I IFN pathway to XP-C pathogenesis but also position this disease within a broader inflammatory framework. Further research is required to fully comprehend the precise mechanisms and implications of this dysregulation in XPC KO keratinocytes and its association with skin inflammatory disorders.

### Conclusion

Xeroderma pigmentosum, complementation group C (XP-C), is a rare autosomal recessive disorder resulting from mutations in the *XPC* gene, leading to impaired nucleotide excision repair and marked sensitivity to ultraviolet radiation. Affected individuals face a significantly elevated risk of early-onset skin cancers and require lifelong photoprotection and vigilant dermatologic monitoring. Despite the absence of curative therapies, understanding the molecular consequences of *XPC* loss remains crucial. This study provides the first evidence of JAK/STAT and type I IFN signaling hyperactivation in XPC KO keratinocytes, offering a remarkable breakthrough into better understanding the molecular mechanisms underlying the disease’s clinical manifestations. Further investigations into the interplay between defects in DNA repair machinery and deregulated inflammatory responses might open new avenues for targeted therapeutic procedures in XPC and other related genetic disorders.

## Materials and Methods

### Cell lines

The human immortalized male epidermal keratinocyte cell line (N/TERT-2G) was generously provided by Dr. James Rheinwald’s laboratory at Harvard Medical School, Boston, USA^15^. The N/TERT-2G cells electroporated with a scrambled sgRNA (negative control) marked as wild type (WT) and the XPC KO cells were cultured in EpiLife medium (60 µM calcium; Gibco™, cat. #MEPI500CA) supplemented with human keratinocyte growth supplement (HKGS, Gibco™, cat. #S0015), containing 0.01 µg/ml recombinant human insulin-like growth factor I, 0.2% (v/v) bovine pituitary extract (BPE), 5 µg/ml bovine transferrin, 0.18 µg/ml hydrocortisone, and 0.2 ng/ml human epidermal growth factor. Additionally, CaCl₂ (340 µM; Sigma-Aldrich, Saint Louis, USA) and 1% penicillin/streptomycin were added. Cells were washed with 8 mL phosphate-buffered saline (PBS, pH 7.4; Gibco™) before dissociation from the T75 cm² culture flask using 3 mL of 0.05% trypsin/EDTA (Gibco™) for 5–10 minutes at 37°C, depending on the cell line. Trypsinization was halted by adding 8 mL of complete culture medium. The cells were then centrifuged at 100× g for 5 minutes, and the supernatant was discarded.

### UVB dose response

The UVB cytotoxicity dataset for N/TERT-2G keratinocytes was previously published in Nasrallah et al., Scientific Reports (2024), doi: 10.1038/s41598-024-81675-6. Here, the 100 J/cm^2^ dose is highlighted and was used as the reference dose for proteomic experiments.

### Kinase phosphosites profiling using PamGene technology

Four samples were prepared in triplicates to decipher the kinase activity signature of wild-type and XPC KO N/TERT-2G keratinocytes at basal state and after UVB irradiation (N=3). The four samples were wild-type non-irradiated, wild type irradiated, XPC KO non-irradiated, and XPC KO irradiated. Cells were seeded in 6 well plates to reach 80% of confluence. Afterwards, they were either irradiated or not and maintained for 1 hour at 37°C. Cells from all experimental conditions were trypsinized and lysed using M-PER™ Mammalian Extraction Buffer (Cat. #78503, Thermo Fisher Scientific, USA) supplemented with Halt™ Protease Inhibitor Cocktail, EDTA-free (100x) (Cat. #78437, Thermo Fisher Scientific, USA) and Halt™ Phosphatase Inhibitor Cocktail (100x) (Cat. #78428, Thermo Fisher Scientific, USA). Protein concentration was determined using the bicinchoninic acid (BCA) or Smith assay, both of which exhibit high sensitivity toward proteins with minimal interference from other substances. Phosphorylation activity was assessed using PamChip® microarrays (PamGene, Cat. #32501), which contain immobilized peptides. Protein samples were prepared by diluting extracted proteins with M-PER buffer supplemented with protease and phosphatase inhibitor cocktails to final concentrations of 5 μg for protein tyrosine kinase assays per condition. The assay protocol followed manufacturer instructions (PamGene International B.V.) as described in the provided protocol documents (https://pamgene.com/wp-content/uploads/2020/09/PTK-assay-protocol-PS12-vs-3-2019-01.pdf). PamChip® microarrays were loaded into the PamStation®12 (PamGene International B.V.), and chips were blocked with 1% bovine serum albumin (BSA) before sample application. A reaction mix containing diluted primary antibodies was added immediately before sample loading. Following a one-and-a-half-hour incubation period, a detection mix was applied to the chips. Fluorescently labeled anti-phospho-antibodies were used to detect phosphorylation activity. The PamStation®12 recorded images of the arrays using a CCD camera. Experiments were performed in triplicate (N=3).

### Kinase phosphosites Profiling: Quality Control and Normalization

A flag system was used to indicate quality control (QC) levels based on three criteria: Phosphosite signal strength (99th percentile of S100 arbitrary units, AU), Phosphosite number control (number of phosphosites passing QC), Sample quality control (coefficient of variance, CV, for technical and biological replicates). All criteria had to be met to receive a green QC flag.

QC levels were determined as follows:

| Level | Flag<sup>1</sup> | Signal Strength (AU) | QC-passed phosphosites (PTK) of 195 | Technical CV | Biological CV |
| --- | --- | --- | --- | --- | --- |
| Good | ● | 0-4,000 | > 123 | < 20% | < 30% |
| Fair | ● | 0-2,000 | 78 - 123 | 20% - 30% | 30% - 40% |
| Poor | ● | 0-1,000 | < 78 | > 30% | > 40% |

Phosphosite QC Selection: For the PTK assay, a kinetic readout was performed with images acquired every 5 minutes. Phosphorylation kinetics were analyzed by plotting S100 values against time, and only phosphosites exhibiting kinetics (signal increase over time) in at least 25% of the arrays (kinetic fraction > 0.25) were included in downstream analysis. Normalization was performed using ComBat correction, which applies empirical Bayes adjustments to reduce systematic bias between PamStation runs and PamChips.

### Proteomic sample preparation

To analyze the proteomic profiles of the electroporated with a scrambled sgRNA (negative control) marked as wild type (WT) and XPC KO N/TERT-2G keratinocytes under basal conditions and after UVB irradiation, four distinct samples were prepared in quadruplicates (N=4): wild-type non-irradiated, wild-type irradiated, XPC KO non-irradiated, and XPC KO irradiated. Cells were cultured in 6-well plates until reaching 80% confluence, then either irradiated or left untreated and incubated for 24 hours at 37°C. Following treatment, cells were trypsinized, and lysed with LYSE-NHS buffer (PreOmics, Germany). Samples were sonicated using a Bioruptor in a 4°C water bath for 10 min, following 30-second cycles with 30 seconds on and 30 seconds off at maximal power. The sample mixture was then centrifuged at 16,000 rpm for 15 minutes at 4°C. The supernatant containing the solubilized proteins was recovered. Protein quantification was performed using either the BCA (bicinchoninic acid) or Smith assay, both of which offer high sensitivity and minimal interference from other substances. Prior to analysis, 10 µg of protein per sample was loaded onto an SDS-PAGE gel to confirm the protein dosage, followed by Coomassie Brilliant Blue staining as per the manufacturer’s instructions (Life Technologies, California, USA). Samples were then prepared using the iST-NHS kit (Preomics), following manufacturer’s instructions. Peptides resulting from LysC/trypsin digestion were labelled using TMTpro™ 16plex Label Reagent Set (ThermoFisher Scientific) before mixing equivalent amounts for further processing. The peptide mix was then fractionated using the Pierce High pH Reversed-Phase Peptide Fractionation Kit (ThermoFisher Scientific).

### LC-MS/MS and proteomic data analysis

The 8 obtained High pH Reversed-Phase fractions were analyzed by online nanoliquid chromatography coupled to MS/MS (Ultimate 3000 RSLCnano and Orbitrap Exploris, Thermo Fisher Scientific) using a 120 min acetonitrile gradient. For this purpose, the peptides were sampled on a precolumn (300 μm x 5 mm PepMap C18, Thermo Scientific) and separated in 75 μm x 250 mm C18 column (Aurora Generation 2, 1.6μm, IonOpticks). The MS and MS/MS data were acquired by Xcalibur (Thermo Fisher Scientific).

Peptides and proteins were identified and quantified using MaxQuant (version 1.6.17.0, PMID: 19029910) with the Uniprot database (*Homo sapiens* taxonomy, 20220322 download), and the frequently observed contaminant database embedded in MaxQuant. Trypsin was chosen as the enzyme and 2 missed cleavages were allowed. Peptide modifications allowed during the search were: C6H11NO (C, fixed), acetyl (Protein N-ter, variable) and oxidation (M, variable). Minimum peptide length and minimum number of razor peptides were respectively set to six amino acids and one. Maximum false discovery rates - calculated by employing a reverse database strategy - were set to 0.01 at peptide and protein levels.

Statistical analysis of MS-based quantitative proteomic data was performed using the ProStaR software (PMID: 27605098). Proteins identified in the reverse and contaminant databases, proteins only identified by site, and proteins quantified in less than four replicates of one condition were discarded. After log2 transformation, extracted corrected reporter abundance values were normalized by Variance Stabilizing Normalization (vsn) method. Statistical testing for comparison of two conditions was conducted with limma, whereby differentially expressed proteins were sorted out using a log2(Fold Change) cut-off of 0.3 and a p-value cut-off of 0.01, leading to a FDR inferior to 1.5 % for each comparison according to the Benjamini-Hochberg estimator.

### Bioinformatic analysis

All data were arranged based on the significance value and fold change so that XPC KO irradiated dysregulated proteins were used as a reference compared to the three other wild-type non-irradiated, wild-type irradiated, and XPC KO non-irradiated to decipher dysregulated proteins either upregulated, downregulated, or unique for XPC KO irradiated cells so that we can unravel their proteomic signature. Several bioinformatics tools were utilized to process the datasets, including StringApp (1.7.0), which makes the visualization and analysis of biological networks better, including all interacting biomolecules. The display of protein-protein interactions can be critically analyzed, and subnetworks can be created for specific interactions and correlations. The String enrichment feature conditions the enrichment analysis and annotates data such as Gene Ontology (GO). Associating the proteome network to different biological functions, such as biological processes, KEGG pathways, Reactome pathways, diseases etc., is instrumental in deciphering the network and bodily functions. FunRich (3.1.3) tool was employed to produce respective heatmaps.

### Immunoblotting

Total proteins were extracted by adding 100 µL of lysis buffer RIPA (Sigma Aldrich, Missouri, USA) supplemented with a phosphatase and protease inhibitor cocktail. A 30-minute incubation of the samples on ice followed this. The sample mixture was transferred to 1.5 mL Eppendorf tubes and centrifuged for 15 minutes at 16000rpm at 4°C. Total protein dosage was further carried out using a BCA protein quantification kit (Life Technologies, California, USA). Western blotting protocol was performed as previously described. Equal protein amounts were resolved by SDS-PAGE (Life Technologies, California, USA) and transferred to a nitrocellulose membrane (IBlot gel transfer, Life Technologies, California, USA). The nitrocellulose membrane was blocked with 5% lyophilized milk or bovine serum album (BSA), followed by the addition of either primary XPC (Santa Cruz 1/500), IFIT-1 (abcam 1/500), IFIT-3 (Santa Cruz 1/500), IRF-9 (Santa Cruz 1/1000), ISG15 (Santa Cruz 1/500), MX1 (abcam 1/500), MX2 (abcam 1/500), PSTAT-1 Tyr701 (Cell Technologies 1/1000), PSTAT-2 Tyr690 (Cell Technologies 1/1000), PSTAT-3 Tyr705 (Cell Technologies 1/1000), antibody incubated overnight at 4°C. Afterwards, incubation with mouse or rabbit HRP antibody (1/5000 or 1/10000 diluted secondary antibody) was done for 1 hour at room temperature and following the addition of the western lightening ECL Pro ECL (Perkin Elmer), images were then directly recorded using Biorad Molecular Imager® Chemi DocTM XRS. Results were analyzed using Image Lab™ software. Glyceraldehyde-3-phosphate dehydrogenase (GAPDH) housekeeping gene was utilized to normalize the expression level of the target gene. Samples were launched in triplicates (N=3).

### Statistical Analysis

GraphPad Prism v.8 was used for statistical analysis, data normalization and quantification of normality to allow the downstream selection of the respective statistical test (parametric or non-parametric) for each particular set of experiments.

## Acknowledgment

AN is supported by a fund from the doctorate school (EDISCE) at University Grenoble Alpes. WR’s contribution was funded by ANR grant PG2HEAL (ANR-18-CE17-0017) and supported by the French National Research Agency in the framework of the “Investissements d’avenir” program (ANR-15-IDEX-02). HR’s and WR’s contribution were funded by ANR grant ANR-23-CE14-0045-01. The proteomic experiments were partially supported by Agence Nationale de la Recherche under projects ProFI (Proteomics French Infrastructure, ANR-10-INBS-08) and GRAL, a program from the Chemistry Biology Health (CBH) Graduate School of University Grenoble Alpes (ANR-17-EURE-0003). The authors thank Dr. Flora Clement for her help with the Pamgene setup.

## Authors’ Contribution

AN performed all the experiments and wrote the manuscript. ES and WR supervised the project. FK and HR revised the manuscript. YC and LB performed MS-based quantitative proteomics. MS, SBV, MFK aided in bioinformatic analysis. All authors edited, read, and approved the manuscript.

## Data availability

All data generated or analysed during this study are included in this published article (and its Supplementary Information files).

